# Dynamic configuration of large-scale cortical networks during an inhibitory task accounts for heterogeneity in attention-deficit/hyperactivity disorder traits

**DOI:** 10.1101/2021.08.04.455077

**Authors:** Jonah Kember, Carolynn Hare, Ayda Tekok-Kilic, William Marshall, Stephen Emrich, Sidney J. Segalowitz, Erin J. Panda

## Abstract

The heterogeneity of attention-deficit/hyperactivity disorder (ADHD) traits (inattention vs. hyperactivity/impulsivity) complicates diagnosis and intervention. Identifying how the configuration of large-scale functional brain networks during cognitive processing correlate with this heterogeneity could help us understand the neural mechanisms altered across ADHD presentations. Here, we recorded high-density EEG while 62 non-clinical participants (ages 18-24; 32 male) underwent an inhibitory control task (Go/No-Go). Functional EEG networks were created using sensors as nodes and across-trial phase-lag index values as edges. Using cross-validated LASSO regression, we examined whether graph-theory metrics applied to both static networks (averaged across time-windows: -500–0ms, 0–500ms) and dynamic networks (temporally layered with 2ms intervals), were associated with hyperactive/impulsive and inattentive traits. Network configuration during response execution/inhibition was associated with hyperactive/impulsive (mean R^2^ across test sets = .20, SE = .02), but not inattentive traits. Post-stimulus results at higher frequencies (Beta, 14-29Hz; Gamma, 30-90Hz) showed the strongest association with hyperactive/impulsive traits, and predominantly reflected less burst-like integration between modules in oscillatory beta networks during execution, and increased integration/small-worldness in oscillatory gamma networks during inhibition. We interpret the beta network results as reflecting weaker integration between specialized pre-frontal and motor systems during motor response preparation, and the gamma results as reflecting a compensatory mechanism used to integrate processing between less functionally specialized networks. This research demonstrates that the neural network mechanisms underlying response execution/inhibition might be associated with hyperactive/impulsive traits, and that dynamic, task-related changes in EEG functional networks may be useful in disentangling ADHD heterogeneity.

## Introduction

Despite being one of the most prevalent behavioral disorders, thought to affect roughly 7.2% of the population under 18 worldwide, the diagnostic reliability of attention-deficit/hyperactivity disorder (ADHD) is an ongoing challenge (1, 2). This is due in part to the heterogeneity of ADHD traits (inattention, hyperactivity, impulsivity), which often fail to neatly align with the distinct presentations recognized by the DSM-V (3). To overcome this, the National Institute of Mental Health proposed that ADHD subtype etiology be refined using biologically based measures, which might better capture the altered mechanisms that account for the distinct traits (4). To date, however, few studies have successfully used biological measures to reliably differentiate the two distinct presentations, characterized by hyperactive/impulsive and inattentive traits (5). The current study examines whether patterns of brain network connectivity recorded from electroencephalogram (EEG) during the execution and inhibition of motor responses can reliably do so.

A primary deficit of those with ADHD is in the ability to appropriately execute and inhibit motor responses based on environmental cues, often operationalized through performance on an inhibitory task (e.g., Go/No-Go, stop-signal, stroop (6, 7, 8)). Dysfunction in the electrocortical responses associated with these functions is consistently implicated in those with ADHD using both fMRI and EEG, although EEG often predicts ADHD with higher accuracy than fMRI (4, 9, 10). In those with ADHD, motor execution/inhibition dysfunction during the Go/No-Go task is evidenced by reduced frontal N2 and central P3 components of the event-related potential (ERP), and decreased event-related oscillatory alpha power (9, 11, 12).

Although inhibitory deficits are pronounced in those with ADHD, some evidence suggests individual differences in the mechanisms underlying the ability to execute/inhibit motor responses might specifically account for hyperactive/impulsive traits (13, 14). In EEG research, for example, compared to the inattentive subtype, those with the combined subtype show higher amplitude beta oscillations at electrodes over the sensorimotor cortex during a cued-flanker inhibitory task (thought to reflect weaker motor response planning (12)). They also show weaker source-localized oscillatory theta power over right frontal areas during the pre-trial intervals of a Go/No-Go task (15). However, contradictory evidence complicates this hypothesis: differences in cognitive measures of inhibitory control often fail to distinguish between subtypes, and electrophysiological research has found no differences between inattentive and hyperactive/impulsive subtypes in the event-related potentials thought to underly motor response inhibition (6, 16, 17, 18, 19).

This controversy seems to suggest that the neural mechanisms supporting motor response execution/inhibition are not a prominent source of heterogeneity in ADHD traits. However, diffusion tensor imaging (DTI) has identified white matter damage to frontal-subcortical circuits and motor circuits (both strongly implicated in response inhibition) as a primary characteristic of the combined subtype compared to the inattentive subtype (20). Additionally, the use of methylphenidate (shown to enhance response inhibition on Go/No-Go tasks by reducing task-irrelevant connectivity), is more effective at reducing hyperactive/impulsive symptoms than inattentive symptoms (21, 22, 23). These results suggest instead that the neural correlates of execution/inhibition are in fact related to ADHD heterogeneity, but that measures capturing neural activity from local regions (MEG/EEG sensors; fMRI regions of interest) might lack the sensitivity required to adequately distinguish between subtypes. Instead, examining patterns of connectivity between cortical regions in the wide-spread functional networks important for cognitive control appears to be necessary. This conclusion– that distinct ADHD symptoms may be linked to altered connectivity between cortical regions during the execution/inhibition of motor responses– has increasingly been made within the broad shift towards understanding ADHD through a network-based approach, rather than a regional-abnormality based approach (5, 14, 24, 25, 26). This network approach is also in line with research examining the neural correlates of response inhibition, which has argued against specific regions functionally specialized for inhibitory processing in favour of a domain-general class of ‘network mechanisms’, where connectivity patterns initiated by the fronto-parietal network account for a variety of functions during cognitive control (27, 28).

One approach recently used in ADHD research is that of task-based network measures, which characterize the dynamic changes in large-scale functional network organization during distinct cognitive processes (14, 29, 30). Since these changes are closely linked to behaviour, it is thought that task-based approaches might better capture sources of ADHD heterogeneity than resting-state approaches (14, 29, 31). A network approach might also further the use of EEG in clinical ADHD research. Indeed, research examining single features in the EEG signal, such as the ratio of theta to beta oscillations or the N2/P3 ERPs, has not yet adequately captured the heterogeneity of ADHD (4, 32, 33). While some research has pursued multivariate approaches to overcome this, where multiple predictive features within the EEG signal are identified using advanced pattern recognition techniques (e.g., convolutional neural networks; 34, 35), their lack of interpretability has so far limited their utility in clinical research as well (4, 34). Thus, the utility of EEG in clinical research may be furthered by research using network measures to identify specific connectivity patterns associated with ADHD heterogeneity (4, 5).

In the current study, we examine whether differences in the organization of functional EEG networks during response execution/inhibition (Go/No-Go task) are associated with distinct ADHD symptoms (inattention and hyperactivity/impulsivity). In line with the Research Domain Criteria (RDoC; dimensional framework proposed by the National Institute of Mental Health), we recognize ADHD traits as lying on a continuum in the broader population (3, 36, 37, 38, 39).

To characterize the dynamic configuration of task-based EEG networks, we use metrics derived from graph theory, which describe the networks as ‘graphs’ (sets of nodes and connections between them, called ‘edges’). Typically, in EEG networks, nodes are sensors, and edges are defined through various functional connectivity measures between sensors. These measures are thought to capture the synchronization of neuronal oscillations from distinct regions, which is a mechanism for neuronal populations to coordinate information transfer (40). In the current study, individual differences in these networks are examined in terms of their integration, segregation, and the balance of these two properties, referred to as ‘small-worldness’. Decreased small-worldness in EEG beta-networks during cognitive interference has been found in those with ADHD compared to controls, and thus might be a specific alteration responsible for ADHD heterogeneity (41). Additionally, we examine functionally specialized collections of nodes called modules, which show strong connectivity with themselves and weak connectivity with the rest of the network (42, 43). Since integration between modules is thought to play a mechanistic role in executive functions (e.g., response inhibition/execution), it may also account for ADHD heterogeneity (43, 44, 45).

Task-based networks are typically analyzed by aggregating over distinct time windows, which conceals important information about how networks may dynamically organize to support response execution/inhibition (46, 47). Here, we examine the utility of a novel dynamic approach, which better captures these changes (46, 47). Given the inherently dynamic way information is transferred across regions during specific cognitive functions, we expected this approach to be well-suited in identifying subtle individual differences in the neural mechanisms supporting response execution/inhibition, and thus be sensitive to differences in ADHD traits.

## Methods and Materials

### Participants

Data were collected from 77 university students. 15 participants were excluded from analysis due to technical problems in data acquisition and a failure to complete the task/questionnaires, resulting in 62 participants entering analysis (see Table 1 for participant info).

**Table 1.** Participant Info and Self-Report Measures.

|  |  |  |
| --- | --- | --- |
| **Demographic Measures** | Age | 20.47 (2.20) |
|  | Sex | 30 Female, 32 Male |
| **Inattentive Measures** | Inattention/Memory | 12.13 (5.61) |
|  | Inattention | 9.77 (4.15) |
| **Hyperactive/Impulsive Measures** | Hyperactivity/Restlessness | 17.89 (6.75) |
|  | Hyperactivity/Impulsivity | 10.06 (4.35) |
|  | Impulsivity/Emotionality | 10.63 (5.38) |

*n* = 62
Results are presented as Mean (SD)

### Conners’ Adult ADHD Rating Scales

Self-report measures of inattention, hyperactivity and impulsivity were collected using Conners’ Adult ADHD Rating scales, which consists of 66 items on a 4-point Likert scale, ranging from “Not at all true” to “Very much true” (48). Five subtests were examined (Table 1), three of which assess hyperactivity and impulsivity, and two of which assess inattention. These scales have high internal and test-retest reliability (α’s = 0.86-0.92; 40).

### Stimuli and Procedure

Participants underwent an A-X continuous performance task (adapted from Tekok-Kilic et al. (49)). Throughout two blocks, 1270 letters (A-H, J, L, X) were presented pseudo-randomly in the center of a computer screen. Participants were asked to respond quickly and accurately with a thumb press on a response pad to the letter sequence ‘A - X’. The letter ‘A’ was presented 200 times, and half of letters following ‘A’ were ‘X’. “Go” trials were when ‘X’ was followed by ‘A’ and required response execution (100 total); “No Go” trials were when another letter followed ‘A’ and required response inhibition (100 total). Each letter was presented for 200ms, and there was an 800ms interval between letters. Mean reaction time and error rates measured task performance. Half of the trials in the task (n = 50) also included a ‘distractor’: simple line-drawings of objects (evenly distributed for Go/No-Go conditions) presented 200ms after ‘A’ for 200ms, although these were excluded from current analyses. This resulted in 50 Go and 50 No-Go trials being used in analysis. During the task, 128-channel EEG was recorded (sampling-rate: 500Hz, impedances kept below 100kΩ) from a HydroCel Geodesic sensor net and 300 NetAmps amplifier (Electrical Geodesics, Inc., Eugene, Oregon). EEG testing took ∼22 minutes, excluding breaks between blocks. Several self-report questionnaires were administered after EEG testing and took ∼20 minutes to complete. For the purposes of the current study, only ADHD scores were analyzed.

### EEG Preprocessing and Phase Synchrony

EEG data were preprocessed in Brain Vision Analyzer 2.2.1 (Brain Products). Data were re-referenced to an average of all sites, and .5Hz high-pass, 100Hz low-pass, and 60Hz notch filters were applied. Gratton and Coles (1983) method was used to correct for eye movements (50), and trials with an amplitude difference of 200μV over a 200ms interval, or with an amplitude >±200μV were rejected, resulting in 42.18 (SD = 5.89) Go trials and 44.35 (SD = 5.41) No-Go trials entering analysis. Connectivity between EEG sensors was calculated as in Panda et al. (51). Specifically, trial-by-trial data (from 3s pre-stimulus to 3s post-stimulus) from each participant were z-scored and filtered into canonical frequency-bands: (1-3Hz; 4-7Hz; 8-13Hz; 14-29Hz; 30-90Hz). Instantaneous phase estimates were obtained using the Hilbert transform, and across-trial phase synchrony was measured using the phase-lag index (PLI; 52). PLI measures the consistency of phase-lags between electrodes across trials but does not imply a directed relationship. As a result, PLI attenuates zero-lag synchrony that might have occurred as a result of volume conduction. With PLI values as edges, sensor-by-sensor adjacency matrices were created at 2ms time-steps for each frequency-band. For dynamic networks, each adjacency matrix was proportionally thresholded to maintain the strongest 10% of edges (resulting in an average of 12.7 edges per node), and binarized. For static networks, adjacency matrices were averaged across time-windows (pre-stimulus/baseline processing: -500-0ms; post-stimulus/task-relevant processing: 0-500ms) before being proportionally thresholded (10%) and binarized.

### Network Analyses

Network analyses were conducted in MATLAB using in-house scripts, the Brain-Connectivity Toolbox (53), and the Dynamic-Graph metrics toolbox (47). Definitions and interpretations of metrics used to characterize the networks are presented in Table 2. These include static and dynamic measures of integration (static: global efficiency; dynamic: broadcast centrality), segregation (static: clustering coefficient; dynamic: temporal correlation coefficient), small-worldness, and modularity (static: modularity, participation coefficient; dynamic: flexibility). Node-level measurements were averaged across all nodes to provide one summary statistic for each participant. For dynamic integration (broadcast centrality), the value halfway between zero and the largest eigenvalue absolute value across all static networks was selected for α (54). To measure how information was transferred between nodes over time, the average ‘burstiness’ of networks was calculated. Communication between nodes is ‘burst-like’ when connections are serially correlated (show random-length periods of sequential connections, followed by random length periods of sequential disconnections), whereas communication is periodic when connections occur at regular intervals (55).

**Table 2.**
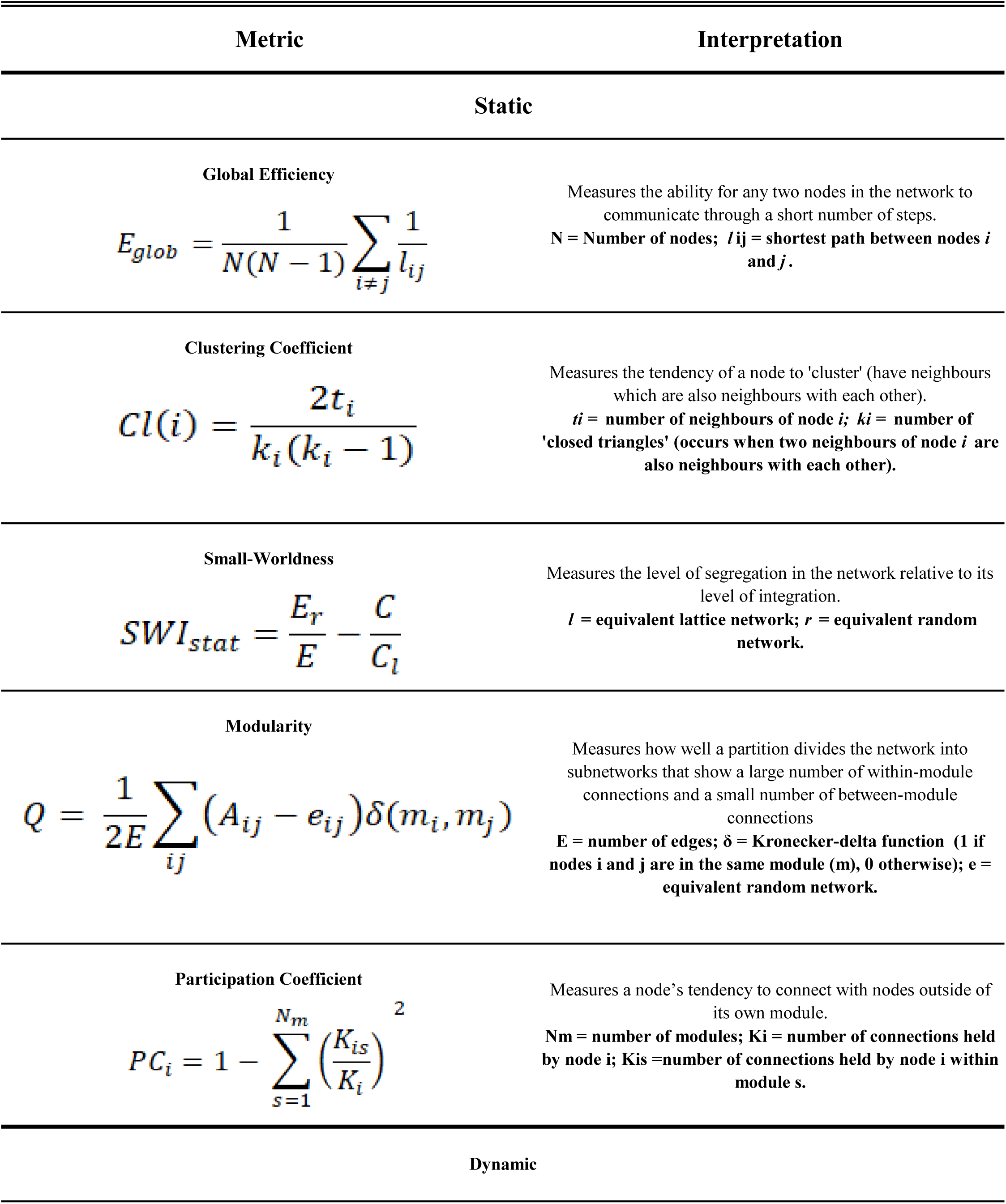
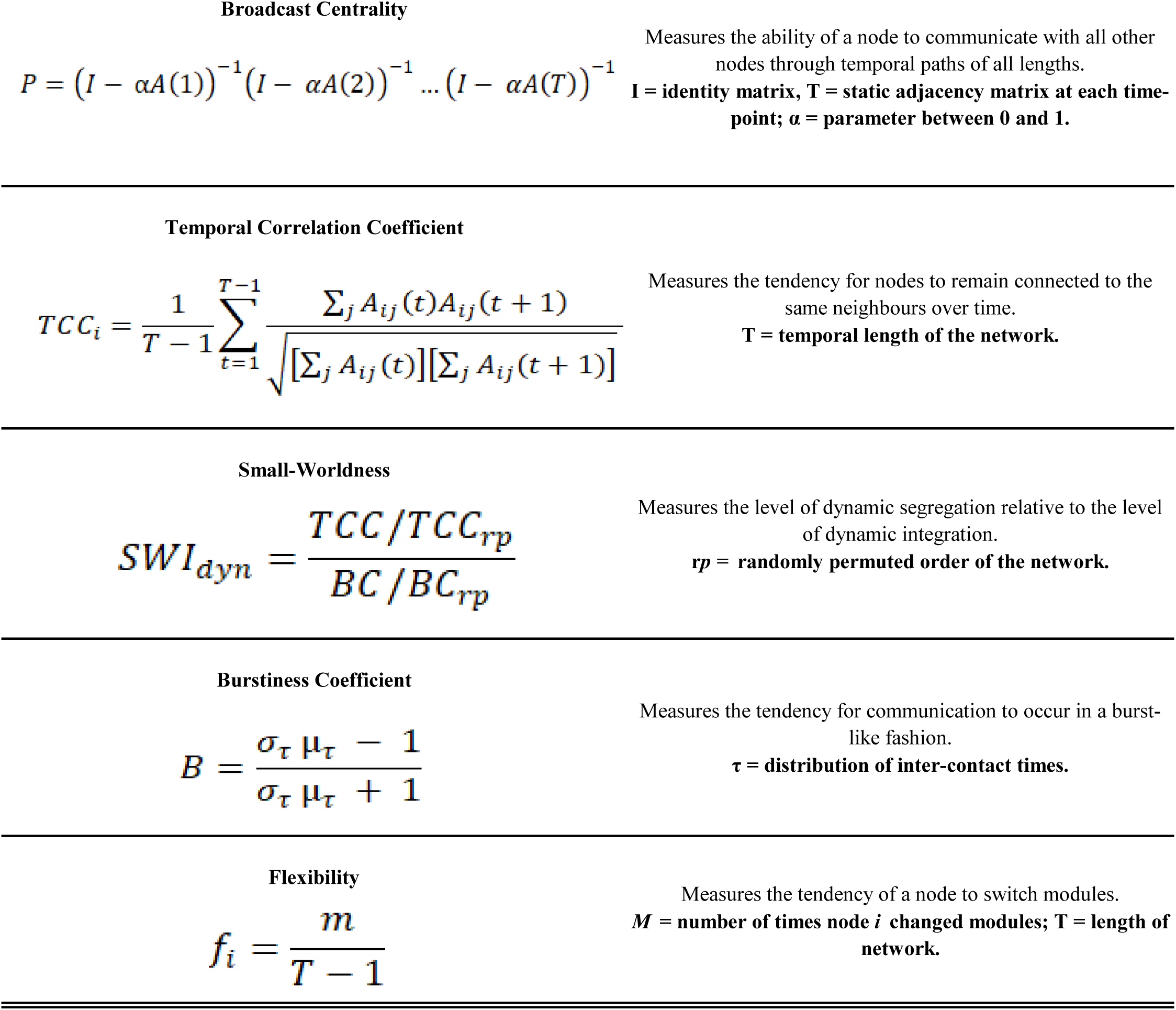
Static and dynamic graph-theory metrics used in analysis.

The optimal partition of each static network into modules was performed using Louvain’s modularity maximization algorithm (γ = 1 (56)). Dynamic modules were detected using the method described by Aynaud & Guillaume (57), which applies the Louvain algorithm to each static network at time *t*, except, rather than the algorithm first assigning each node to their own module, nodes are first assigned to the module with which they belonged at *t* – 1. This avoids issues that arise from community-detection methods which identify modules at each timepoint independently (58). Due to the small variance that arises from Louvain’s algorithm, flexibility was averaged across 100 runs (59).

### Regression Analyses

To identify network characteristics (features) associated with hyperactivity/impulsivity and inattention (outcome variables), regression analyses were conducted using the least absolute shrinkage and selection operator (LASSO; 60). LASSO is a statistical learning method that fits data to a multiple linear regression and prevents overfitting by applying a penalty term (which includes a hyperparameter lambda) to the cost function used to estimate regression coefficients. When lambda is zero, the LASSO simplifies to the least-squares estimates; when lambda is sufficiently high, all regression coefficients become zero. By driving small coefficients to zero, LASSO results in a relatively interpretable model with a sparse set of features. The optimal lambda value, which maximizes the model’s ability to account for variability in future observations, was estimated using repeated 10-fold cross-validation. This process is as follows: 1) partition the data into 10 equal folds, 2) fit the model to all but one of these folds using a wide range of lambda values (training set; normalized using the min-max method), 3) for each lambda value, measure the mean squared error (MSE) of the model on the fold left out (test set), 4) calculate the out-of-sample R^2^ on this test set using the lambda value that minimizes MSE, 5) using each fold as a test set, determine the average MSE across test sets as a function of lambda, 6) estimate regression coefficients using the lambda value that minimizes this average MSE. Ten repetitions of this k-fold cross validation were used to determine MSE as a function of lambda before selecting the appropriate value (as this helps avoid biases of unrepresentative partitions). This lambda value was used to estimate a final model on the entire dataset (providing the regression coefficients to be interpreted). To assess the quality of the model, the average out-of-sample R^2^ across test sets was used. To interpret the model, regression coefficients and Pearson’s *r* were used.

## Results

### AX-CPT Task Performance

Across participants, mean reaction time was 360.2 ms (*SD* = 79.5) and an average of 0.15 (*SD* = 0.36) commission errors (incorrect response on No-Go trials) were made. The most commission errors made by a participant was four; most participants (n = 38) made zero.

### LASSO Models

Separate regression models were estimated for the five ADHD scales during both the pre- and post-stimulus windows (−500–0ms; 0–500ms). Static and temporal network measures from all frequency-bands and both Go/No-Go conditions were model features. Models conducted on post-stimulus data (0-500ms) predicted measures of hyperactivity and impulsivity (Hyperactivity/Impulsivity: R^2^ = .202 SE = .020; Hyperactivity/Restlessness: R^2^ = .188, SE = .022; Impulsivity/Emotionality: R^2^ = .123, SE = .023; average R^2^ across test sets provided). This can be seen in Figure 1. Conversely, models for inattentive traits had all regression coefficients driven to zero (Inattention: R2 = .033, SE = .025; Inattention-Memory: R2 = .029, SE = .020), suggesting the relationship between inattention and network configuration was not strong enough to overcome the overfitting penalty. All models conducted on data from the pre-stimulus period (−500–0ms) had regression coefficients driven to zero as well, suggesting baseline network configuration did not relate to ADHD traits (Inattention: R2 = .051, SE = .017; Inattention/Memory: R2 = .071, SE = .014; Hyperactivity/Restlessness: R2 = .031, SE = .046; Hyperactivity/Impulsivity: R2 = .082, SE = .015; Impulsivity/Emotionality: R2 = .078, SE = .022).

**Figure 1.**
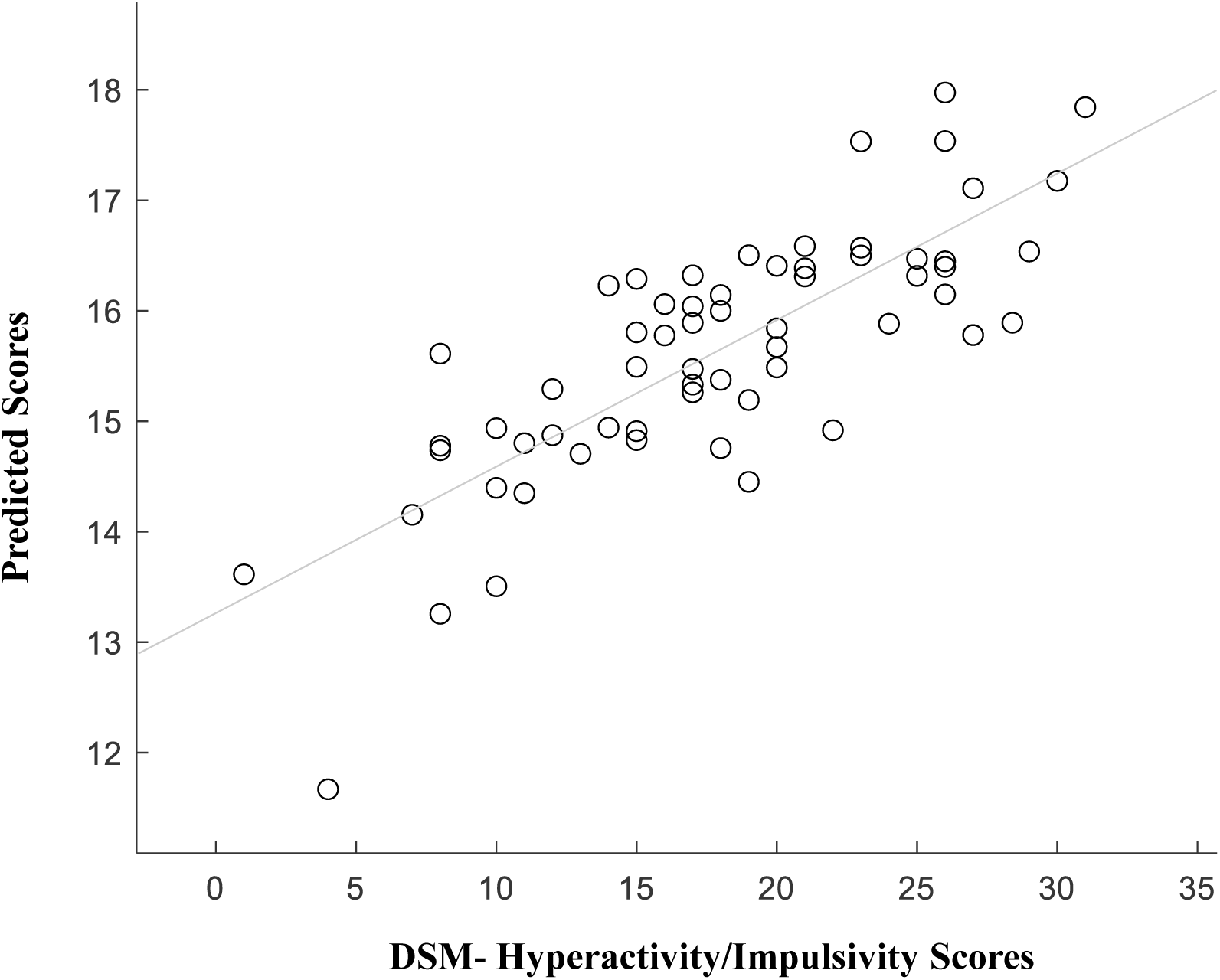
Empirical and predicted hyperactive/impulsive behaviours. Scatterplot of empirical and predicted scores (from LASSO model) on Hyperactivity/Impulsivity (least squares regression line plotted in gray). Model included all network measures in low and high frequency bands for both Go and NoGo conditions in the post-stimulus condition. Across all test-sets used in cross validation, the average out of sample R^2^ was .202.

### Condition Analyses

Features identified in the Go and No-Go conditions were then examined separately to understand differences between the execution (Table 3) and inhibition of motor responses (Table 4). Results are presented for Hyperactivity/Impulsivity (R^2^ = .202) and Hyperactivity/Restlessness (R^2^ = .188).

**Table 3.** Execution of a motor response (Go)

**Hyperactivity/Impulsivity**
| Features |  |  | Strength |  |  |
| --- | --- | --- | --- | --- | --- |
| Freq | Class | Measure | $\beta$ | Pearson's $r$ | 95% CI |
| Beta | Static | Integration | -2.48 | - |  |
| Beta | Static | Modularity | 1.41 | .30, $p = .0178$ | [.06 .51] |
| Beta | Dynamic | Burstiness | -1.24 | -.32, $p = .0167$ | [-.52 -.06] |
| Beta | Static | Participation Coefficient | -1.11 | -.34, $p = .0064$ | [-.55 -.10] |
| Beta | Dynamic | Integration | -.90 | -.22, $p = .089$ | [-.44 .03] |
| Alpha | Static | Modularity | .57 | - |  |
| Gamma | Dynamic | Flexibility | .17 | .25, $p = .0482$ | [.01 .47] |

**Hyperactivity/Restlessness**
| Features |  |  | Strength |  |  |
| --- | --- | --- | --- | --- | --- |
| Freq | Class | Measure | $\beta$ | Pearson's $r$ | 95% CI |
| Beta | Static | Participation Coefficient | -2.14 | -.28, $p = .023$ | [-.49 -.03] |
| Beta | Static | Integration | -1.91 | - |  |
| Beta | Dynamic | Integration | -.41 | - |  |
| Gamma | Dynamic | Small-Worldness | 3.95 | 0.40, $p = .0011$ | [.17 .59] |
| Gamma | Static | Integration | -2.06 | -0.23, $p = .076$ | [-.45 .02] |
| Alpha | Static | Segregation | 3.10 | - |  |

Features in the LASSO models which were identified as predictive of scores on the CAARS scales. In the model (one run for each scale), all frequency bands, static and dynamic measures, as well as Go and NoGo conditions were features. For each feature in the Go condition, regression coefficients ( $\beta$ ), pearson's $r$ (when $p > .1$ ), and 95% confidence intervals for pearson's $r$ are provided.

**Table 4.** Inhibition of a motor response (No-Go)

**Hyperactivity/Impulsivity: No-Go**
| Features |  |  | Strength |  |  |
| --- | --- | --- | --- | --- | --- |
| Freq | Class | Measure | $\beta$ | Pearson's $r$ | 95% CI |
| Gamma | Static | Small-Worldness | 4.76 | .43, $p = .0006$ | [.20 .61] |
| Gamma | Dynamic | Integration | 2.70 | 0.36, $p = .0038$ | [.12 .56] |
| Gamma | Dynamic | Small-Worldness | 2.16 | - |  |
| Gamma | Static | Integration | 1.16 | .31, $p = .01$ | [.06 .52] |
| Gamma | Dynamic | Segregation | -.44 | -.36, $p = .0043$ | [-.56 -.12] |
| Alpha | Dynamic | Integration | 0.32 | - |  |
| Theta | Static | Modularity | 0.31 | - |  |

**Hyperactivity/Restlessness: No-Go**
| Features |  |  | Strength |  |  |
| --- | --- | --- | --- | --- | --- |
| Freq | Class | Measure | $\beta$ | Pearson's $r$ | 95% CI |
| Gamma | Static | Small-Worldness | 5.60 | .41, $p = .0008$ | [.18 .60] |
| Gamma | Static | Integration | 4.90 | 0.33, $p = .01$ | [.082 .53] |
| Gamma | Static | Participation Coefficient | 2.13 | .35, $p = .0055$ | [.11 .55] |
| Beta | Dynamic | Integration | -1.86 | -.28, $p = .029$ | [-.49 -.03] |
| Delta | Dynamic | Integration | -.49 | -.21, $p = .09$ | [-.43 .04] |

Features in the LASSO models which were identified as predictive of scores on the CAARS scales. In the model (one run for each scale), all frequency bands, static and dynamic measures, as well as Go and NoGo conditions were features. For each feature in the No-Go condition, regression coefficients ( $\beta$ ), pearson's $r$ (when $p > .1$ ), and 95% confidence intervals for pearson's $r$ are provided.

In the Go condition, hyperactive/impulsive traits were associated with the configuration of EEG networks following stimulus presentation (0–500ms). The majority of network features (Hyperactivity/Impulsivity: 5/7, Hyperactivity/Restlessness: 3/6) were in the beta-band (14-29 Hz; regression coefficients and Pearson’s *r* presented in Table 3). The results in Table 3 tell us that during response executive in those with high hyperactivity/impulsivity, EEG networks oscillating at a beta frequency become less integrated, show a more modular structure, tend to form connections within these modules, and communicate more periodically. This dynamic configuration can be seen in Figure 2, and predominantly reflects less long-range/burst-like communication between frontal-central and bilateral-posterior regions around 200–300ms in those with high hyperactivity/impulsivity.

**Figure 2.**
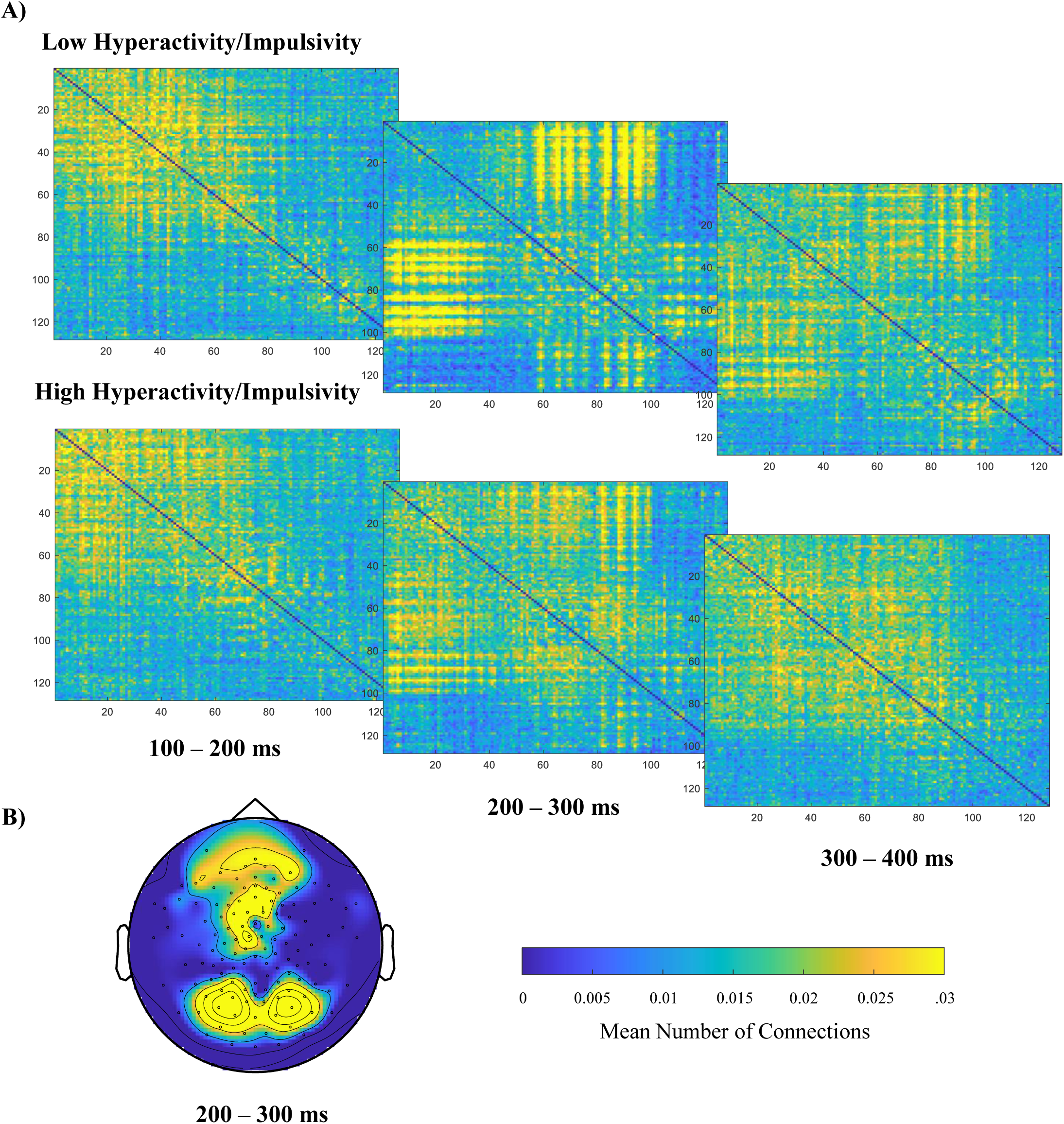
Post-stimulus beta network configuration (14-29Hz) during execution of a response (Go) in those with high and low hyperactivity/impulsivity. (A) Adjacency matrices with sensors as nodes, and the mean number of connections within each time-window as edges. Network are averaged across the 10 participants with the lowest and highest hyperactivity scores. (B) Topographical map showing the mean number of connections from 200-300ms (thresholded at 10% and binarized), averaged across all participants (*n* = 62).

Similarly, in the No-Go condition, hyperactive/impulsive traits were associated with the configuration of high-frequency EEG networks following stimulus presentation, although the majority of features (Hyperactivity/Impulsivity: 5/7, Hyperactivity/Restlessness: 3/5) were in the gamma networks (30-90Hz; Table 4). The results in Table 4 tell us that networks oscillating at a gamma-frequency in those with high hyperactivity/impulsivity are less stable over time, less segregated, and more integrated. However, they exhibit a level of segregation higher than expected based on their level of integration (small-worldness), suggesting the configuration did not simply tend towards randomness. This configuration can be seen in Figure 3, and predominantly reflects communication within left frontal-central regions that is less segregated in those with high hyperactivity/impulsivity.

**Figure 3.**
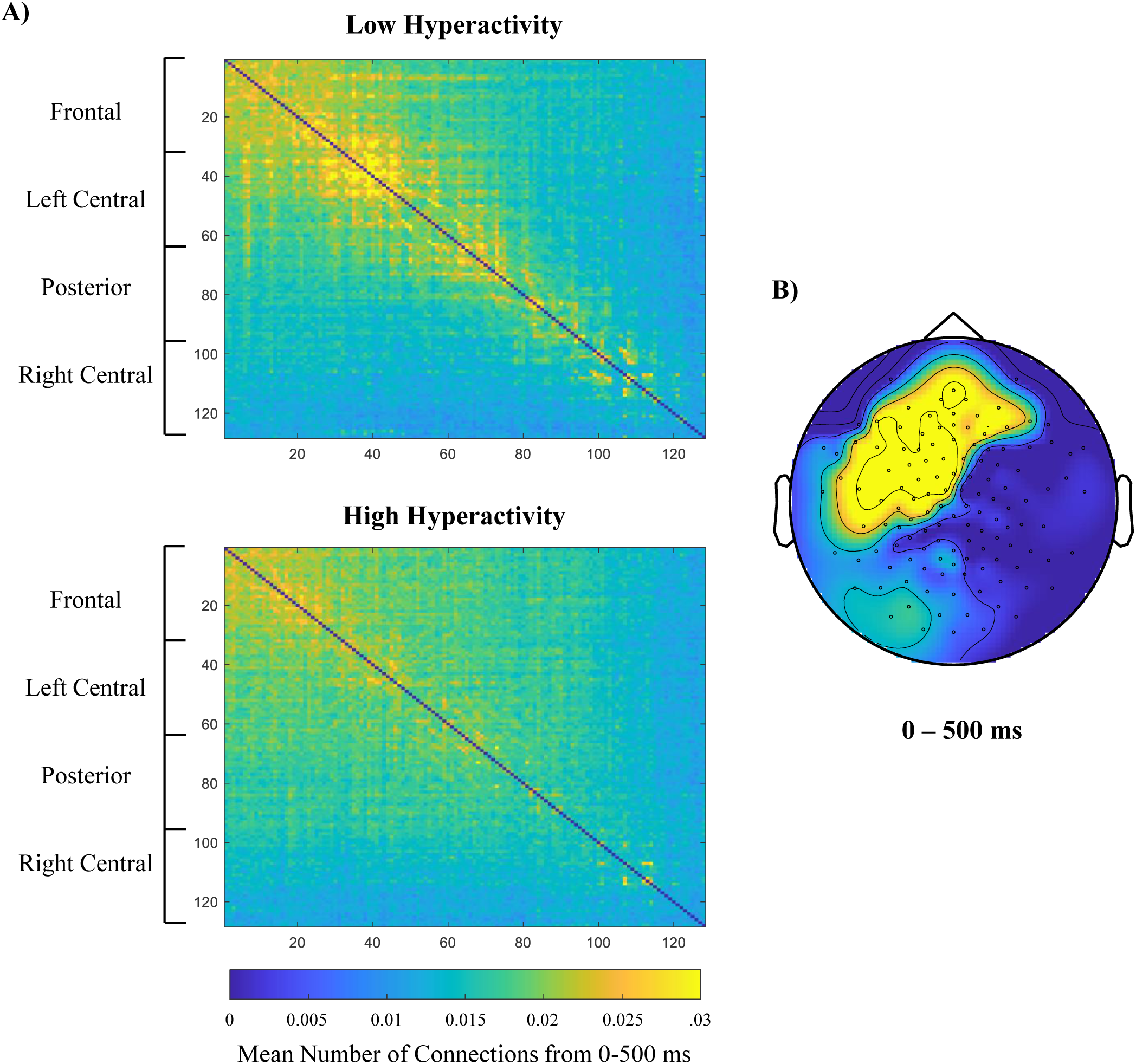
Post-stimulus (0-500ms) gamma network (30-90Hz) configuration during inhibition of a response (NoGo) in those with high and low hyperactivity/impulsivity. (A) Adjacency matrices with sensors as nodes, and the mean number of connections from 0-500ms as edges. Each network is averaged across the 10 participants with the lowest and highest hyperactivity scores. (B) Topographical map with the mean number of connections show by each node over the same time period (thresholded at 10% and binarized).

## Discussion

In this study, we investigated whether the dynamic configuration of large-scale cortical networks during the execution and inhibition of motor responses acts as a source of heterogeneity in ADHD. We found that the dynamic configuration of EEG networks during both response execution (Go) and inhibition (No-Go) is altered in those with hyperactive/impulsive, but not inattentive, traits. Specifically, hyperactivity/impulsivity was linked to decreased/less burst-like integration in networks oscillating at a beta frequency during response execution, and increased integration/small-worldness in networks oscillating at a gamma frequency during response inhibition. These results suggest that differences in how functional networks are dynamically recruited during response execution/inhibition may contribute to ADHD heterogeneity.

### Motor Response Execution: Weaker Beta Network Integration Associated with Hyperactivity/Impulsivity

During response execution (Go), networks oscillating at a beta frequency showed a more modular configuration (i.e., could easily be divided into subnetworks), with less integration between modules as hyperactivity/impulsivity increased. Dynamically, these networks lacked the burstiness (clear on/off periods) seen in those with less hyperactivity/impulsivity. Cortical beta oscillations have long been implicated in motor control, and bursts of beta communication within the motor cortex are thought to reflect the appropriate selection and initiation of movements (61, 62). When considered alongside the location of electrodes involved (bilateral central-posterior and frontal-central) and its timing (∼200-300ms), this finding might reflect difficulty initiating integration of prefrontal and motor circuit activity (see Figure 2). This is similar, although posterior, to the increased oscillatory beta power found over central-bilateral motor areas during response preparation in the combined versus inattentive subtype (12). Our results further explain this effect by suggesting it arises from alterations to the mechanisms driving integration between prefrontal and motor circuitry.

Given the structure and myelination of white matter tracts is thought to strongly constrain EEG connectivity (63), this interpretation may be consistent with abnormal white matter structure previously reported in the combined ADHD subtype (20, 64). Abnormalities in the circuits associated with motor control/inhibition (supplementary motor area and middle frontal gyrus (20)) would be particularly supportive of this explanation and would suggest the lack of beta integration observed in those with hyperactive/impulsive traits arises from white matter tract abnormalities that hinder the integration between specialized frontal and motor systems.

This interpretation, however, should be made with caution. First, difficulty suppressing default mode network (DMN) activity during response execution could also influence the integration and modularity of oscillatory beta activity (DMN interference hypothesis: 65, 66, 67, 68). Second, there is some evidence that the DTI findings thought to characterize the combined ADHD subtype (20) might not hold up to more stringent analyses practiced today (69). Future research, then, might benefit from investigating how white matter abnormalities indexed by radial/axial diffusivity (19, 62) restrict the electrophysiological mechanisms driving integration between neural circuits (using, for example, dynamic tractography techniques, which examine the relationship between cortico-cortical evoked potentials and diffusion MRI measures (70)).

### Motor Response Inhibition: Stronger Gamma Network Integration/Small-Worldness associated with Hyperactivity/Impulsivity

During response inhibition (No-Go), hyperactivity/impulsivity was associated with altered short-range gamma connectivity within left frontal-central areas. Specifically, with greater hyperactivity/impulsivity, gamma networks became more integrated/small-world like, showed more between-module communication and were more likely to change over time. This is in line with previous findings of decreased oscillatory frontal-gamma power from 300-600ms for Go/No-Go conditions in those with ADHD versus non-ADHD controls (9). Event-related fNIRS research has suggested reduced activity during Go/No-Go tasks in ADHD might be localized to circuitry in the left prefrontal cortex (71).

While the “Go” beta network configuration may reflect a mechanism that is insufficiently evoked (an inability to integrate functionally specialized systems in the prefrontal and motor cortex), this “No-Go” gamma network configuration may instead reflect a mechanism evoked to a greater degree. First, increased integration, small-worldness, and between-module processing in gamma networks is the same configuration observed during increased cognitive load (e.g., to increased *n* in n-back working memory tasks (72)). These changes are thought to occur when specialized sub-networks within the cortex coordinate their processing to successfully complete the required function (59, 72). Second, increased small-worldness, a configuration thought to minimize metabolic costs while maximizing the potential for complex interactions, was seen in the gamma networks of those with hyperactive/impulsive traits (73, 74, 75). Rather than a shift towards random or regular structure (an alteration commonly seen in the functional networks of those with psychiatric disorders), increased small-worldness is thought to be advantageous during periods of increased cognitive demand (73, 74, 75). Together, these results suggest that those with hyperactive/impulsive traits respond to the task as though it is more cognitively demanding and requires a higher level of integration/small-worldness to be properly completed. In other words, those with hyperactive/impulsive traits may have less functionally specialized networks and evoke an integrative mechanism to compensate (76).

### Lack of Association Between Inattention and Configuration of Alpha Networks

To our surprise, we found little association between the configuration of oscillatory alpha networks and ADHD traits. In attentional networks during Go/No-Go tasks, alpha oscillations are thought to gate information transfer between fronto-parietal and occipital regions, thereby suppressing task-irrelevant and highlighting task-relevant sensory information (77, 78, 79). A large body of research examining electrophysiological correlates of ADHD has suggested individual differences in this alpha may explain inattentive symptoms (80, 81). However, in this study, network configuration did not predict inattentive traits, and alpha results for hyperactive/impulsive traits were inconsistent (small regression coefficients and no linear correlations). The relative simplicity and consistent inter-stimulus interval of the current task might have limited the extent to which attentional gating varies within typically developing adults. Future research measuring EEG network configuration during more cognitively demanding tasks (i.e., cued reaction-time task) might highlight altered mechanisms found within the inattentive subtype.

### Limitations

Our findings should be considered alongside certain limitations. First, we used a conservative estimate of connectivity (phase-lag index), which attenuates ‘pure’ (zero-lag) synchronization in an attempt to account for volume conduction (52). Since some of the attenuated synchronization likely reflects true neural connectivity between distinct regions rather than volume conduction effects (82), methodological advances that allow for the interpretation of pure EEG synchronization (i.e., inverse modelling advances) will allow us to draw conclusions about these mechanisms more confidently. Second, while this research benefitted from a dimensional approach, the extent to which ADHD behaviours are dimensional versus categorical is poorly understood, and consideration through both lenses appears to be required (83). Because of this, the extent to which those with clinical levels of hyperactivity/impulsivity will exhibit the same alterations is unclear. We predict similar findings (weaker beta integration/stronger gamma integration), albeit more pronounced.

## Conclusion

Accurately distinguishing inattentive and hyperactive/impulsive presentations is important when deciding on the appropriate intervention. This is evidenced, for example, by the reduced response to methylphenidate found in the inattentive subtype (23). By demonstrating that the neural mechanisms underlying this heterogeneity can be captured through functional network measures applied to the EEG recorded during a Go/No-Go task, this research furthers the use of EEG in clinical research focused on delineating the ADHD subtypes.

## Acknowledgments

We would like to extend our thanks to those who took the time to participate in the study, and to Abraham Omorogieva for his help with data collection.

